# Symbolic Kinetic Models in Python (SKiMpy): Intuitive modeling of large-scale biological kinetic models

**DOI:** 10.1101/2022.01.17.476618

**Authors:** Daniel R. Weilandt, Pierre Salvy, Maria Masid, Georgios Fengos, Robin Denhardt-Erikson, Zhaleh Hosseini, Vassily Hatzimanikatis

## Abstract

**Motivation:** Large-scale kinetic models are an invaluable tool to understand the dynamic and adaptive responses of biological systems. The development and application of these models have been limited by the availability of computational tools to build and analyze large-scale models efficiently. The toolbox presented here provides the means to implement, parametrize and analyze large-scale kinetic models intuitively and efficiently.

**Results:** We present a Python package (SKiMpy) bridging this gap by implementing an efficient kinetic modeling toolbox for the semiautomatic generation and analysis of large-scale kinetic models for various biological domains such as signaling, gene expression, and metabolism. Furthermore, we demonstrate how this toolbox is used to parameterize kinetic models around a steady-state reference efficiently. Finally, we show how SKiMpy can imple-ment multispecies bioreactor simulations to assess biotechnological processes.

**Availability:** The software is available as a Python 3 package on GitHub: https://github.com/EPFL-LCSB/SKiMpy, along with adequate documentation.

## 1 Introduction

Organisms are complex and adaptive systems, posing a challenge when investigating their response to environmental or genetic perturbations (Kitano, 2002). In this context, large-scale kinetic models are an essential tool to understand how the underlying biochemical reaction networks respond to such perturbations (Chowdhury *et al*., 2015). However, currently, no modeling framework allows users to build and analyze large-scale kinetic models efficiently. Therefore, we propose a novel Python toolbox that enables the user to semiautomatically reconstruct a kinetic model from a constraint-based model (Salvy *et al*., 2019). Furthermore, we express the models in terms of symbolic expressions, allowing the straightforward implementation of various analysis methods, e.g., numerical integration of the ordinary differential equations (ODEs).

Such numerical analysis requires a set of kinetic parameters describing the individual reaction characteristics. However, as parameters from literature or databases (Schomburg *et al*., 2013) are collected *in vitro* and often fail to capture the *in vivo* reaction kinetics (Weilandt and Hatzimanikatis, 2019), a series of methods have been developed to infer parameters from phenotypic observations (Khodayari and Maranas, 2016; Saa and Nielsen, 2016; Gonzalez *et al*., 2007; Wang *et al*., 2004). To this end, we here present the first open-source implementation of the ORACLE framework to efficiently generate steady-state consistent parameter sets (Wang *et al*., 2004; Miskovic and Hatzimanikatis, 2010; Chakrabarti *et al*., 2013; Savoglidis *et al*., 2016; Tokic *et al*., 2020).

## 2 Methods

### Implementing kinetic models

The system of ordinary differential equations describing the kinetics of a biochemical reaction network can be derived directly from the mass balances of the *N* reactants participating in the *M* reactions of the network:

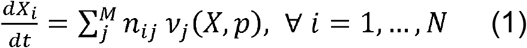

where *X*_*i*_ denotes the concentration of the chemical *i, n*_*ij*_ is the stoichiometric coefficient of reactant *i* in reaction *j* and *v*_*j*_ (***X, p***) is the reaction rate of reaction *j* as a function of the concentration state variables ***X*** = [*X*_1_, *X*_2_,…, *X*_*N*_ ]^*T*^ and *K* kinetic parameters ***p*** = *p*_1_, *p*_2_,…, *p*_*K*_]^*T*^. The functions *v*_*j*_ (***X, p***) are the given rate laws of each reaction *j*. An overview of the implemented rate laws is given in Table S1. Are the reactants distributed across multiple compartments of each reactant’s mass balance is modified according to (For details, see supplementary material):

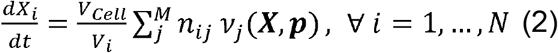

where *V*_*Cell*_ is the overall cell volume and *V*_*i*_ is the compartment volume for concentration *X*_*i*_.

### Efficient steady-state consistent parametrization

To overcome the scarcity of kinetic data, SKiMpy provides the means to infer the parameters efficiently on a large scale by sampling sets of kinetic parameters consistent with steady-state physiology (Wang *et al*., 2004; Wang and Hatzimanikatis, 2006; Miskovic and Hatzimanikatis, 2010). These parameter sets are then evaluated with respect to local stability, global stability, and relaxation time to discard unstable models and models with non-physiological dynamics.

## 3 Usage

This toolbox implements various methods and resources that allow the user to build and analyze large-scale kinetic models in an efficient manner with a detailed account of the implemented methods given in the supplemental material. We further provide different tutorials demonstrating the modeling capabilities (Fig 1 A-C, for details, see supplementary material)

**Fig. 1.**
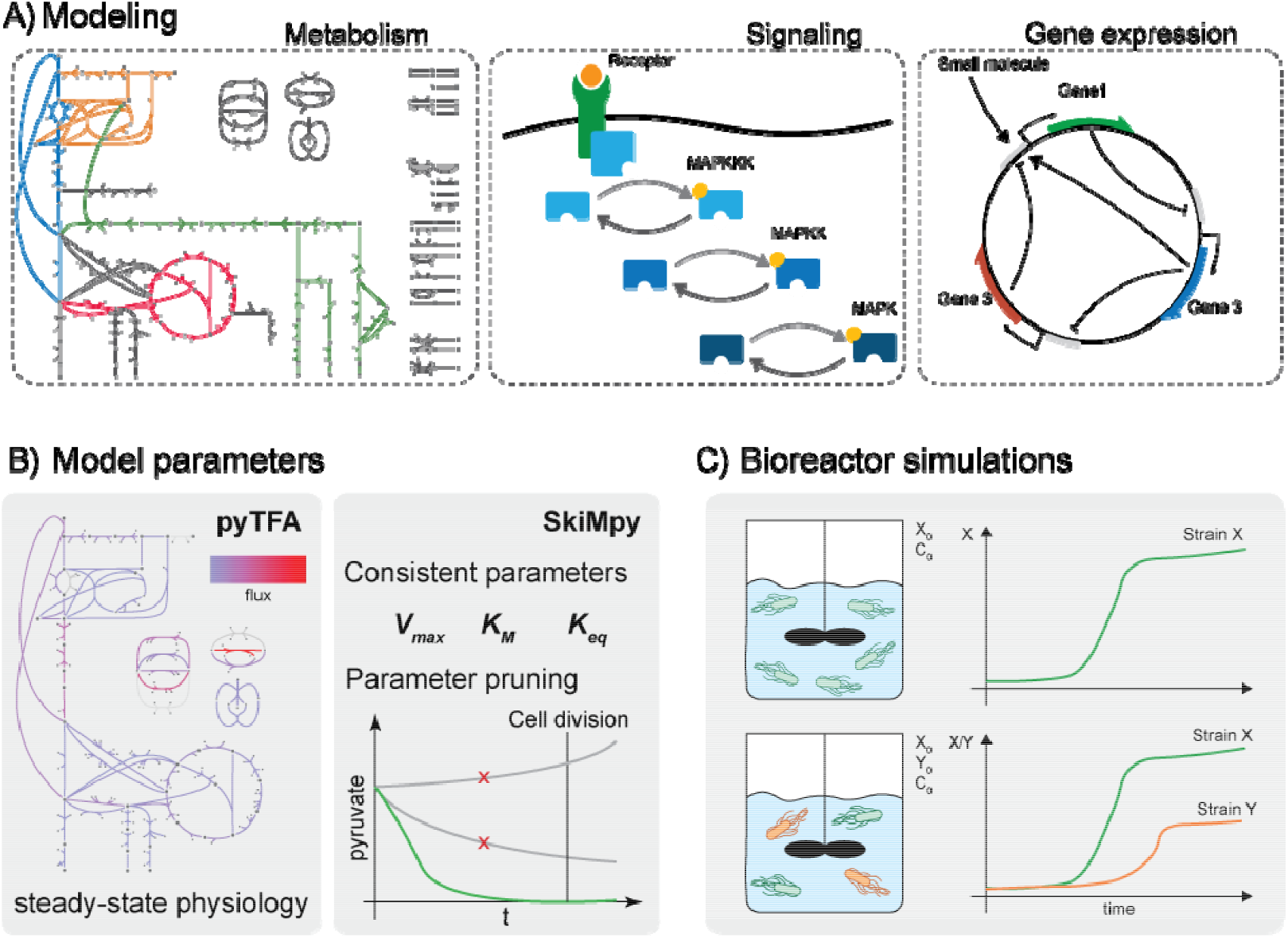
Software capabilities. A) Building and simulation of different types of models B) parameterization of kinetic metabolic models around a reference steady-state C) reactor modeling and multispecies simulations

## 4 Conclusion

SKiMpy enables the user to reconstruct kinetic models for large-scale biochemical reaction systems. With an extensive palette of analytic methods, the software presents a versatile platform to model biological systems, as shown with the examples given in the supplemental material with models of i) *E. coli’*s metabolism, ii) a signaling pathway, iii) synthetic gene-expression circuits, and iv) different strains in a bioreactor. Furthermore, as the software generates symbolic expressions directly available to the user, SKiMpy facilitates the implementation of novel parameter inference and consistent sampling methods. Thus, SKiMpy represents a method development platform to analyze cell dynamics and physiology on a large scale. Finally, the presented toolbox increases the accessibility of large-scale kinetic models to various biological disciplines and studies ranging from biotechnology to the medical sciences.

## Acknowledgments

The authors want to thank Ljubisa Miskovic for valuable discussions.

## Funding

This work has received funding from the European Union’s Horizon 2020 Research and Innovation Programme under the Marie Skłodowska-Curie grant agreements No. 722287 and 675585, the European Union’s Horizon 2020 Research and Innovation Programme under grant agreement No. 686070 and 814408 as well as by the Swiss National Centre of Competence in Research (NCCR) Microbiomes.

## Conflict of Interest

none declared.

## Notes

https://github.com/EPFL-LCSB/SKiMpy

### Competing Interest Statement

The authors have declared no competing interest.

