## Supplementary information for "Symbolic Kinetic Models in Python (SKiMpy): Intuitive modeling of large-scale biological kinetic models"

#### Supplementary introduction

This toolbox implements various methods and resources that allow the user to build and analyze large scale kinetic models efficiently, including i) automated draft model construction from FBA and TFA using Cobrapy[1] and pyTFA[2], respectively, ii) an extensive library of kinetic mechanisms, iii) efficient model parametrization using ORACLE [3–7], iv) local stability analysis[8], v) global stability analysis, vi) local sensitivity analysis, e.g., metabolic control analysis[9], vii) global sensitivity analysis and uncertainty propagation[10,11] viii) modal analysis[8], iix) non-linear ODE integration[12,13], ix) identification of conserved pools[14]. Therefore, the presented Python package implements an object-oriented interface to construct the symbolic expressions from a library of kinetic mechanisms using sympy[15] and then precompiling these expressions into machine code using Cython [16]. SKiMpy also integrates the SUNDIALS ODE-Solver package [13] using the interface provided by the ODES package[17].

This supplementary material provides additional information on the capabilities and use-cases of the SKiMpy toolbox. We first introduce the general workflow (Figure S1) and then give a detailed overview of the library of implemented kinetic mechanisms. We then introduce the concept of kinetic modifiers and their relation to the kinetic mechanism. Subsequently, we demonstrate the modeling and analysis capabilities of the SKiMpy toolbox for three examples from different biological domains, i.e., gene circuits, signaling, and metabolism. Therefore, we implement previously published models of the mitogen activate kinase pathway[18] and stripe-forming synthetic gene circuits[19].

Additionally, we created a model of *E. coli*'s central carbon metabolism with genome-scale biosynthetic requirements based on the metabolic model by Varma et al.[20] and the genome-scale metabolic model (GEM) iJO1366[21] using the ORACLE workflow[3,4,22] as implemented in SKiMpy (Supplementary Material and Methods). Furthermore, we show how the toolbox SKiMpy can be used to simulate organism-specific models in a bioreactor. Finally, we introduce how the toolbox allows implementing global sensitivity for metabolic control analysis[10,11] using the ORACLE framework.

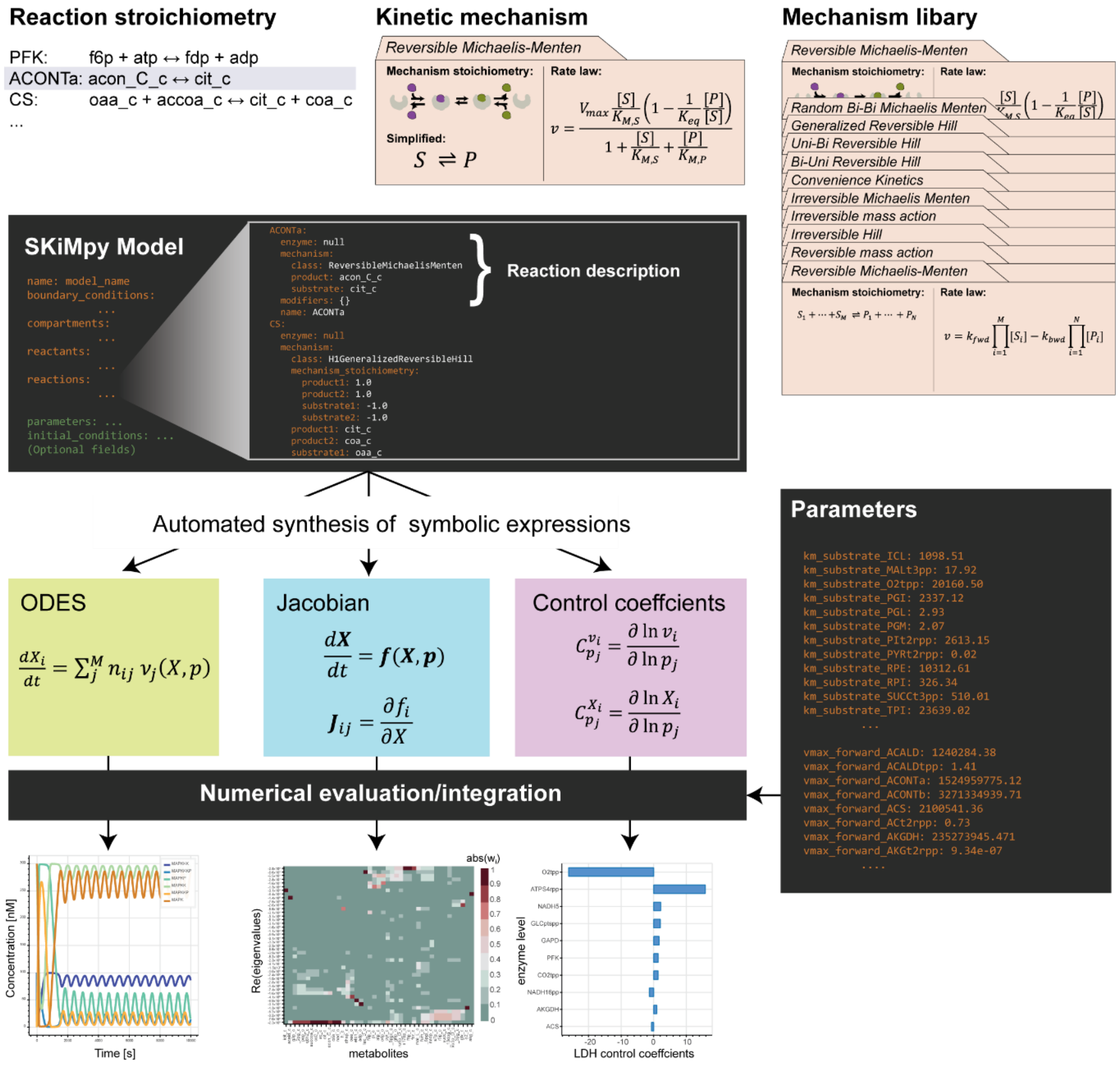


**Figure S1:** SKiMpy modeling workflow: Kinetic models are built using consistent kinetic mechanisms from a library of kinetic mechanisms. Saving the SKiMpy model in the YAML format allows the user to edit the reaction descriptions intuitively. The generic SKiMpy model can then be used to compile the different kinds of symbolic expressions describing, for example, i) ordinary differential expressions of the system, ii) the system's Jacobian, or iii) the control coefficients. Given a set of parameters, SKiMpy further directly allows to evaluate or integrate these expressions numerically.

#### General Formulation of non-linear kinetic models with compartments

The chemical reactions governing the cellular functions can occur in different compartments[23]. Hence, a single molecule yields a different concentration in each compartment depending on the respective volume of this compartment. It is thus necessary to account for the size of the respective compartments to adjust the changes of concentration upon molecule transport. We derive this relation from the mass balance:

$$\frac{d\boldsymbol{m_x}}{dt} = N\boldsymbol{v_m}$$

where $\boldsymbol{m_x}$ are the molecule numbers $\boldsymbol{v_m}$ are the mass fluxes, and $N$ is the stoichiometric matrix. Reformulation in terms of concentrations yields:

$$\frac{d(V\boldsymbol{x})}{dt} = \frac{dV}{dt}\boldsymbol{x} + V\frac{d\boldsymbol{x}}{dt} = N\boldsymbol{v_m}$$

$$V^{-1}\frac{dV}{dt}\boldsymbol{x} + \frac{d\boldsymbol{x}}{dt} = V^{-1}N\boldsymbol{v_m}$$

$$\frac{d\boldsymbol{x}}{dt} = V^{-1}N\boldsymbol{v_m} - \mu\boldsymbol{x}$$

where $V$ is a diagonal matrix with the respective compartment volume $V_i$ for each compound $X_i$. It has been shown that for metabolic reaction systems, the dilution due to cell growth $\mu\boldsymbol{x}$ is small compared to the reaction rates and can therefore be neglected[24]. Hence, the system can be described in terms of its mass fluxes and concentration as:

$$\frac{d\boldsymbol{x}}{dt} = V^{-1}N\boldsymbol{v_m}$$

Typically it is common to report reaction rates in [M]/[s] instead of mass fluxes [mol]/[s] when describing reaction kinetics. As reaction rates have units of concentration per time, they require an associated reference volume. Considering $V_{ref}$ as a diagonal matrix with the reference volume $V_{ref,j}$ for every reaction rate $v_j$ then,

$$\frac{d\boldsymbol{x}}{dt} = V^{-1}NV_{ref}\boldsymbol{v}$$

When modeling the reactions using elementary steps, this reference volume is typically the volume in which the collision between the reactants yields the respective reaction. For transport reactions, the units of the forward and backward rate and rate constants differ, making it difficult to compare parameters in large-scale kinetic models. In SKiMpy, we overcome these heterogeneous units by scaling all reaction rates to units of the cell volume $V_{cell}$. Therefore, the ODE system can be expressed according to:

$$\frac{dX_i}{dt} = \frac{V_{Cell}}{V_i} \sum_{j}^{M} n_{ij} \, v_j(\boldsymbol{X}, \boldsymbol{p}),$$

This way, the $V_{max}$ and $k_{f/b}$ parameters of the rate law $v_j(\boldsymbol{X}, \boldsymbol{p})$ retain consistent units and can directly be compared with each other.

#### Kinetic mechanisms

The SKiMpy toolbox aims to implement a large variety of Kinetic mechanisms and behaviors. We, therefore, provide an extensive list of explicit and generic mechanisms (Table S1). To increase the modeling capabilities and allow additional model refinement, we further implemented the concepts of reaction flux modifiers (Table S2). These modifiers allow the user to implement additional kinetic effects on the forward and reverse flux expressions increasing the modeling repertoire combinatorically. For example, these modifiers can be used to model the effect of small molecules on the reaction kinetics in case these do not have an explicit binding site on the enzyme, e.g., protons. We, therefore, developed two kinds of modifiers i) the first-order small molecule modifier, which assumes that the small molecules directly affect the $k_{cat}$ in the form of a mass-action relationship and ii) the displacement small molecule modifier modeling a case where the small molecules are assumed to affect the enzyme kinetics other than in $k_{cat}$. For the latter case, we know from the Wegscheider condition[25] that the small molecule concentrations will affect the effective equilibrium constant $K_{eq}$ (Table S2).

**Table S1:** Kinetic mechanisms implemented in SKiMpy

| Name | Reaction | Rate law repression |
| --- | --- | --- |
| Michaelis Menten | $S \rightleftharpoons P$ | $v = \frac{V_{max} \frac{[S]}{K_{M,S}} \left(1 - \frac{1}{K_{eq}} \frac{[P]}{[S]}\right)}{1 + \frac{[S]}{K_{M,S}} + \frac{[P]}{K_{M,P}}}$<br>Parameters: $[V_{max}, K_{M,S}, K_{M,P}, K_{eq}]$ |
| Random Bi-Bi<br>Michaelis Menten | $S_1 + S_2 \rightleftharpoons P_1 + P_2$ | $v = \frac{V_{max} \frac{[S_1]}{K_{M,S_1}} \frac{[S_2]}{K_{M,S_2}} \left(1 - \frac{1}{K_{eq}} \frac{[P_1][P_2]}{[S_1][S_2]}\right)}{1 + \frac{[S_1]}{K_{M,S_1}} + \frac{[S_2]}{K_{M,S_2}} + \frac{[P_1]}{K_{M,P_1}} + \frac{[P_2]}{K_{M,P_2}} + \frac{[S_1][S_2]}{K_{M,S_1} K_{I,S_2}} + \frac{[P_1][P_2]}{K_{M,P_2} K_{I,P_1}}}$<br>Parameters: $[V_{max}, K_{M,S_1}, K_{M,S_2}, K_{I,S_2} K_{M,P_1}, K_{M,P_2}, K_{I,P_1} K_{eq}]$ |
| Generalized<br>Reversible Hill[26] | $S_1 + \cdots + S_N \rightleftharpoons P_1 + \cdots + P_N$ | $v = \frac{V_{max} \left(1 - \frac{1}{K_{eq}} \prod_{i=1}^{N} \left(\frac{[P_i]}{[S_i]}\right)\right) \prod_{i=1}^{N} \left(\frac{[S_i]}{K_{M,S_i}}\right) \left(\frac{[S_i]}{K_{M,S_i}} + \frac{[P_i]}{K_{M,P_i}}\right)^{h-1}}{\prod_{i=1}^{N} \left(1 + \left(\frac{[S_i]}{K_{M,S_i}} + \frac{[P_i]}{K_{M,P_i}}\right)^{h}\right)}$<br>Parameters: $[V_{max}, K_{M,S_1} \ldots K_{M,S_M}, K_{M,P_1} \ldots K_{M,P_M}, K_{eq}, h]$ |
| Uni-Uni Reversible<br>Hill with modifiers[26] | $\downarrow/\perp$<br>$S_1 \rightleftharpoons P_1$ | $v = \frac{V_{max} \left(\frac{[S_1]}{K_{M,S_1}}\right) \left(1 - \frac{1}{K_{eq}} \frac{[P_1]}{[S_1]}\right) \left(\frac{[S_1]}{K_{M,S_1}} + \frac{[P_1]}{K_{M,P_1}}\right)^{h-1}}{\prod_{j}^{N_m} \left[\frac{\left(1 + \left(\frac{[M_j]}{[K_{M,j}]}\right)^{h}\right)}{\left(1 + \sigma_j^{2h} \left(\frac{[M_j]}{[K_{M,j}]}\right)^{h}\right)}\right] + \left(\frac{[S_1]}{K_{M,S_1}} + \frac{[P_1]}{K_{M,P_1}}\right)^{h}}$<br>Parameters: $[V_{max}, K_{M,S_1}, K_{M,P_1}, K_{eq}, h, K_{M,1}, .. K_{M,j}, \sigma_1, \ldots, \sigma_j]$ |
| Bi-Bi Reversible<br>Hill with modifiers[26] | $\downarrow/\perp$<br>$S_1 + S_2 \rightleftharpoons P_1 + P_2$ | |

| | | |
|---|---|---|
| | | $v = \frac{V_{max}\left(\frac{[S_1]}{K_{M,S_1}}\frac{[S_2]}{K_{M,S_2}}\right)\left(1-\frac{1}{K_{eq}}\frac{[P_1][P_2]}{[S_1][S_2]}\right)\left(\frac{[S_1]}{K_{M,S_1}}+\frac{[P_1]}{K_{M,P_1}}\right)^{h-1}\left(\frac{[S_2]}{K_{M,S_2}}+\frac{[P_2]}{K_{M,P_2}}\right)^{h-1}}{\Pi_j^{N_m}\left[\frac{\left(1+\left(\frac{[M_j]}{[K_{M,j}]}\right)^h\right)}{\left(1+\sigma_j^{4h}\left(\frac{[M_j]}{[K_{M,j}]}\right)^h\right)}\right]+\Pi_j^{N_m}\left[\frac{\left(1+\sigma_j^{2h}\left(\frac{[M_j]}{[K_{M,j}]}\right)^h\right)}{\left(1+\sigma_j^{4h}\left(\frac{[M_j]}{[K_{M,j}]}\right)^h\right)}\right]\left[\left(\frac{[S_1]}{K_{M,S_1}}+\frac{[P_1]}{K_{M,P_1}}\right)^h\left(\frac{[S_2]}{K_{M,S_2}}+\frac{[P_2]}{K_{M,P_2}}\right)^h\right]^{N_m}\left(\frac{[S_1]}{K_{M,S_1}}+\frac{[P_1]}{K_{M,P_1}}\right)^h\left(\frac{[S_2]}{K_{M,S_2}}+\frac{[P_2]}{K_{M,P_2}}\right)^h}$<br>Parameters: $[V_{max}, K_{M,S_1}. K_{M,S_2}, K_{M,P_1}, K_{M,P_2}, K_{eq}, h, K_{M,1}, .. K_{M,j}, \sigma_1, \dots, \sigma_j]$ |
| Uni-Bi Reversible Hill[26] | $S_1 \rightleftharpoons P_1 + P_2$ | $v = \frac{V_{max}\left(\frac{[S_1]}{K_{M,S_1}}\right)\left(1-\frac{1}{K_{eq}}\frac{[P_1][P_2]}{[S_1]}\right)\left(\frac{[S_1]}{K_{M,S_1}}+\frac{[P_1]}{K_{M,P_1}}\frac{[P_2]}{K_{M,P_2}}\right)^{h-1}}{1+\left(\frac{[S_1]}{K_{M,S_1}}+\frac{[P_1]}{K_{M,P_1}}\right)^h+\left(\frac{[S_1]}{K_{M,S_1}}+\frac{[P_2]}{K_{M,P_2}}\right)^h+\left(\frac{[S_1]}{K_{M,S_1}}+\frac{[P_1]}{K_{M,P\_1}}+\frac{[P_2]}{K_{M,P_2}}\right)^h-2\left(\frac{[S_1]}{K_{M,S_1}}\right)^h}$<br>Parameters: $[V_{max}, K_{M,S_1}, K_{M,P_1}, K_{M,P_2}, K_{eq}, h]$ |
| Bi-Uni Reversible Hill[26] | $S_1 + S_2 \rightleftharpoons P_1$ | $v = \frac{V_{max}\left(\frac{[S_1]}{K_{M,S_1}}\frac{[S_2]}{K_{M,S_2}}\right)\left(1-\frac{1}{K_{eq}}\frac{[P_1]}{[S_1][S_2]}\right)\left(\frac{[S_1]}{K_{M,S_1}}\frac{[S_2]}{K_{M,S_2}}+\frac{[P_1]}{K_{M,P_1}}\right)^{h-1}}{1+\left(\frac{[S_1]}{K_{M,S_1}}+\frac{[P_1]}{K_{M,P_1}}\right)^h+\left(\frac{[S_2]}{K_{M,S_2}}+\frac{[P_1]}{K_{M,P_1}}\right)^h+\left(\frac{[S_1]}{K_{M,S_1}}+\frac{[S_2]}{K_{M,S_2}}+\frac{[P_1]}{K_{M,P\_1}}\right)^h-2\left(\frac{[P_1]}{K_{M,P_1}}\right)^h}$<br>Parameters: $[V_{max}, K_{M,S_1}, K_{M,S_2}, K_{M,P_1}, K_{eq}, h]$ |
| Convenience Kinetics[27] | $\alpha_1 S_1 + \cdots + \alpha_M S_M \rightarrow \beta_1 P_1 + \cdots + \beta_N P_N$ | $v = \frac{V_{max}\Pi_{i=1}^{M}\left(\frac{[S_i]}{K_{M,S_i}}\right)\left(1-\frac{1}{K_{eq}}\frac{\Pi_{j=1}^{N}[P_j]}{\Pi_{i=1}^{M}[S_i]}\right)}{\Pi_{i=1}^{M}\Sigma_{m=0}^{\alpha_i}\left(\frac{[S_i]}{K_{M,S_i}}\right)^m+\Pi_{j=1}^{N}\Sigma_{m=0}^{\beta_j}\left(\frac{[P_j]}{K_{M,P_j}}\right)^m-1}$<br>Parameters: $[V_{max}, K_{M,S_1} \dots K_{M,S_M}, K_{M,P_1} \dots K_{M,P_N}, K_{eq},]$ |
| Irreversible Michaelis Menten | $S_1 + \cdots + S_M \rightarrow P_1 + \cdots + P_N$ | $v = V_{max}\prod_{i=1}^{M}\left(\frac{\frac{[S_i]}{K_{M,S_i}}}{1+\frac{[S_i]}{K_{M,S_i}}}\right)$<br>Parameters: $[V_{max}, K_{M,S_0} \dots K_{M,S_M}]$ |
| Reversible mass action | $S_1 + \cdots + S_M \rightleftharpoons P_1 + \cdots + P_N$ | $v = k_{fwd}\prod_{i=1}^{M}[S_i] - k_{bwd}\prod_{i=1}^{N}[P_i]$<br>Parameters: $[k_{fwd}, k_{bwd}]$ |
| Irreversible mass action | $S_1 + \cdots + S_M \rightarrow P_1 + \cdots + P_N$ | $v = k_{fwd}\prod_{i=1}^{M}[S_i]$<br>Parameters: $[k_{fwd},]$ |
| Irreversible Hill | $S \rightarrow P$ | $v = V_{max}\frac{\left(\frac{[S]}{K_{M,S}}\right)^h}{1+\left(\frac{[S]}{K_{M,S}}\right)^h}$<br>Parameters: $[V_{max}, K_{M,S}, h]$ |
| Approximate gene expression with regulation[19] | 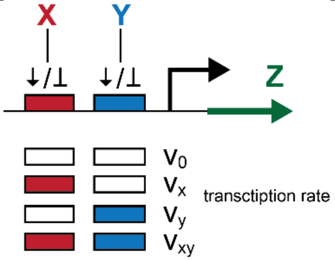 | $v_z = \frac{v_0 + v_x\left(\frac{[X]}{K_X}\right)^n + v_y\left(\frac{[Y]}{K_y}\right)^m + v_{xy}c\left(\frac{[X]}{K_X}\right)^n\left(\frac{[Y]}{K_y}\right)^m}{1+\left(\frac{[X]}{K_X}\right)^n+\left(\frac{[Y]}{K_y}\right)^m+c\left(\frac{[X]}{K_X}\right)^n\left(\frac{[Y]}{K_y}\right)^m}$<br>Parameters: $[v_0, v_x, v_y, v_{xy}, K_X, K_y, c, n, m]$ |

**Table S2:** Kinetic modifiers implemented in SKiMpy, $v = v_f - v_r$ denotes the mechanism net flux where $v_f$ is the forward and $v_r$ is the reverse flux expression. Tilda denotes the respective modified fluxes.

| Name | Effect | Rate law modification |
|---|---|---|
| Simple Inhibitor | | $\tilde{v} = \frac{v}{\left(1+\frac{[I]}{K_I}\right)}$<br>Parameters: $[K_I]$ |
| Simple Activator | | $\tilde{v} = v\left(1+\frac{[A]}{K_A}\right)$<br>Parameters: $[K_A]$ |

| | | |
|---|---|---|
| First-order small molecule modifier | $n_s C_s + S \rightarrow P$<br>$S \rightarrow P + n_p C_p$ | $\tilde{v}_f = v_f [C]^{n_s}$<br>$\tilde{v}_r = v_b [C]^{n_p}$<br>$\tilde{v} = \tilde{v}_f - \tilde{v}_r$ |
| Displacement small molecule modifier | $n_s C_s + S \rightarrow P$<br>$S \rightarrow P + n_p C_p$ | $\tilde{v}_f = v_f$<br>$\tilde{v}_r = v_b \frac{[C]^{n_p}}{[C]^{n_s}}$<br>$\tilde{v} = \tilde{v}_f - \tilde{v}_r$ |

### Kinetic model implementation

This section uses SKiMpy to implement kinetic models for a prototypical singling system, a genetic circuit, and a metabolic reaction network. All these examples can be found in tutorials within the software package.

#### Example signaling network

To demonstrate the capability of SKiMpy to build and simulate kinetic models of signaling pathways, we build a prototypical signaling cascade of the Mitogen-activated protein kinases[18] (Figure S1a). This model consists of input in the form of a receptor kinase activity that phosphorylates the Mitogen-activated protein kinase kinase kinases (MAKKK). The phosphorylated form of MAKKK (MAKKKP) then phosphorylates the Mitogen-activated protein kinase kinases (MAPKK), which in their phosphorylated form (MAPKKP) phosphorylate the Mitogen-activated protein kinases (MAPK), which acts as the output signal of this system. A dephosphorylating phosphatase counters each phosphorylation reaction. We model the kinase and phosphatases using the irreversible Michaelis-Menten Kinetics (Table S1). The inhibition kinetics are implemented via the simple inhibitor mechanism (Table S2). We parametrize the resulting models with parameters similar to the ones found by Kholodenko et al.[18]. The resulting models show a Hill like steady-state response in the output concentrations for increasing input activities (Figure S1b) and oscillatory behavior for the case when feedback inhibition is active.

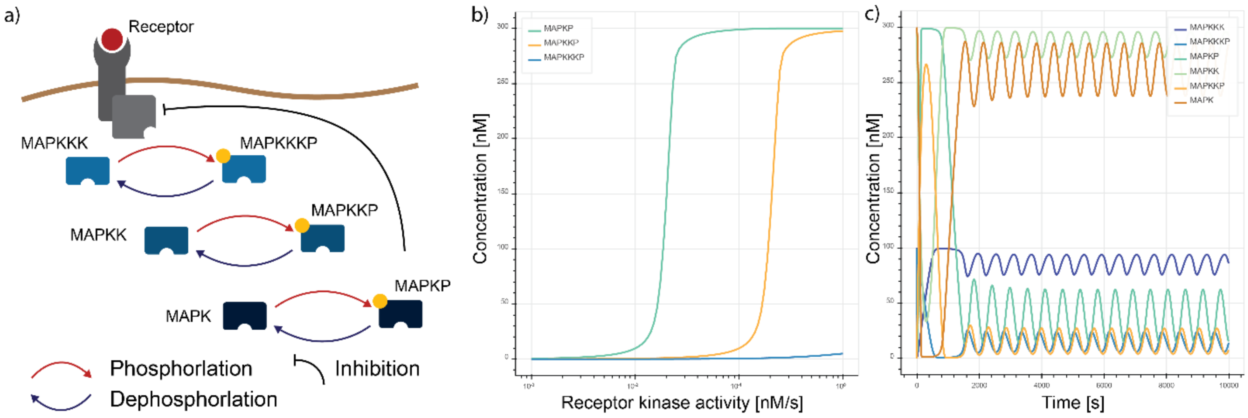


**Figure S2**: a) Signaling cascade model, b) Steady-State output for different receptor inputs without feedback inhibition, c) Oscillatory behavior with feedback activation.

#### Example Genetic Circuit

In the next step, we implement a stripe forming genetic circuit from the collection of genetic circuits presented by Schaerli et al.[19] (Figure S2a). We implement this kinetic model using a slightly generalized version of the approximate transcription kinetics used by Schaerli et al. (Table S1) combined with irreversible mass action kinetics for the mRNA degradation. Using the parameters from the supplementary material of Schaerli et al.[19], we are able to reproduce one of the stripe-forming phenotypes (Figure S2b).

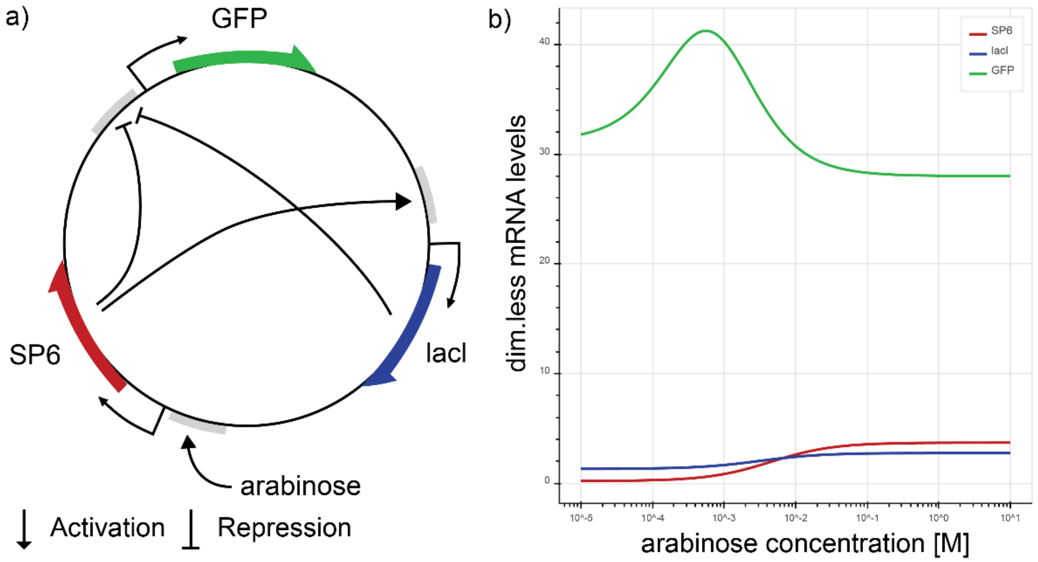


**Figure S3**: a) Schematic representation of the genetic circuit, b) Steady-State response of expression level to different arabinose concentrations.

#### Example Metabolic model

Finally, we demonstrate the implementation of metabolic kinetic models with SKiMpy using a model of the central carbon metabolism of *E. coli* (Figure S6). This kinetic model was built and parameterized around a reference steady-state (Figure S3a) based on a stoichiometric model using the ORACLE workflow. The details of the kinetic model and the ORACLE workflow can be found in the Supplementary Materials and Methods. Here we use this model to demonstrate how modal analysis, metabolic control analysis, and large parameter perturbations can be used to investigate large-scale kinetic models.

Using a numerical formulation based on the reaction rates elasticities, SKiMpy can efficiently compute the Jacobian of the non-linear system at the steady-state[28,29,3]. The eigenvalues and eigenvectors of the Jacobian can be used to analyze the dynamic behavior for small perturbations from a steady-state via Modal analysis[8]. Thereby the complex contribution $w_{ij}$ of each concentration $[X_j]$ in the system to a pool variable $y_i$ is calculated as:

$$y_i = \sum_{i}^{N} w_{ij} [X_j]$$

Each of these pool variables $y_i$ (also called modes) follows the dynamics of a complex exponential function:

$$y_i(t) = e^{\lambda_i t}$$

where $\lambda_i$ is the respective complex eigenvalue. If the real part of all eigenvalues is smaller than zero, the deviation from the steady-state vanishes over time, and the system returns to the steady-state such systems are called locally stable. Using the ORACLE framework ensures that all models are locally stable at the reference steady-state (see below).

The weights $w_{ij}$ allow understanding how much a concentration $[X_j]$ contributes to the mode $y_i$. If a variable has a large absolute contribution $|w_{ij}|$ to a specific mode $y_i$ this variable will follow the exponential decay of the respective eigenvalue $\lambda_i$. Plotting and clustering these contributions in a matrix format (Figure S3b) allows identifying pools consisting of similar concentrations and concentrations that appear with high contributions in the same modes. Such an analysis also identifies fast and slow-moving variables[30].

SKiMpy further implements the numerical log(linear) formulation[3,22,28,29] to calculate metabolic control coefficients for fluxes and concentrations. These control coefficients allow accessing how fluxes and concentration will change for small perturbations. We calculated the flux control coefficients of lactate dehydrogenase (LDH) for all enzyme concentrations and ranked them according to their absolute value (Figure S3c). As the first three supposed targets are transport reactions, we selected the test the design of the fourth and fifth targets, namely increasing the enzyme concentration of LDH and the Glyceraldehyde-3-phosphate dehydrogenase (GAPD).

Finally, we use the full set of non-linear equations to integrate the ODEs with the reference steady-state as the initial conditions. We then implement the selected design and overexpress LDH and GAPD two-fold. Monitoring the lactate flux as a function of time shows that a two-fold increase in GAPD results in a 20% increase of the LDH flux, whereas a two-fold increase of LDH leads to a decrease of the LDH flux.

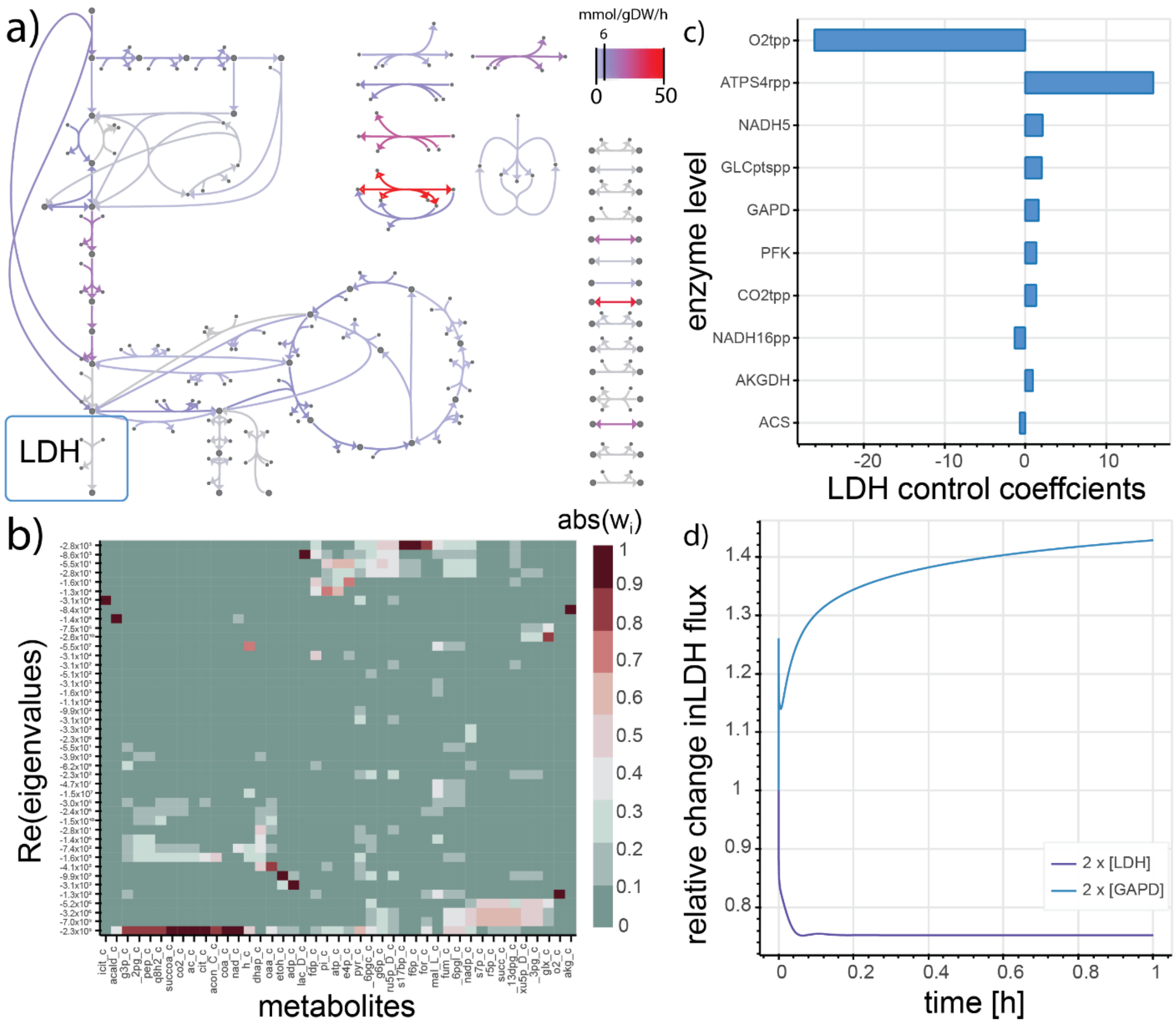


**Figure S4**: a) Escher map of the metabolic reaction network colored by the steady-state flux b) Absolute value of the modal matrix of the metabolic models allow clustering the dynamics around the steady-state c) control coefficients for the production of lactate via the lactate dehydrogenase reaction (LDH), d) Temporal responses of the lactate production flux upon a two-fold increase of the LDH and the Glyceraldehyde-3-phosphate dehydrogenase (GAPD) enzyme concentrations respectively.

#### Reactor simulations

When simulating large-scale kinetic models, the environment is often assumed to be invariant over time. This assumption holds only for short observations times. For longer observation times, it is necessary to observe how the concentrations in the environment change as a function of time. To address this issue, we implemented the concept of reactors. These reactors allow modeling the experimental environment and run simulations with one or multiple species as different kinetic models.

To demonstrate these capabilities of SKiMpy, we construct a batch reactor model. For a typical bioproduction process with a single *E. coli* strain. We inoculate a 1L reactor with about 0.1g/L of cells and 20g/L of glucose. We assume the oxygen, CO2, and proton concentration to be constant over the process. The results show that our kinetic model can capture the exponential growth phase until glucose deletion and then growth arrests (Figure S4a/b).

We repeat the simulation and add an *E. coli* with the GAPD overexpression to increase the lactate production as presented above. The mutant exhibits a slightly higher growth rate than the wild-type (Figure S4c/d).

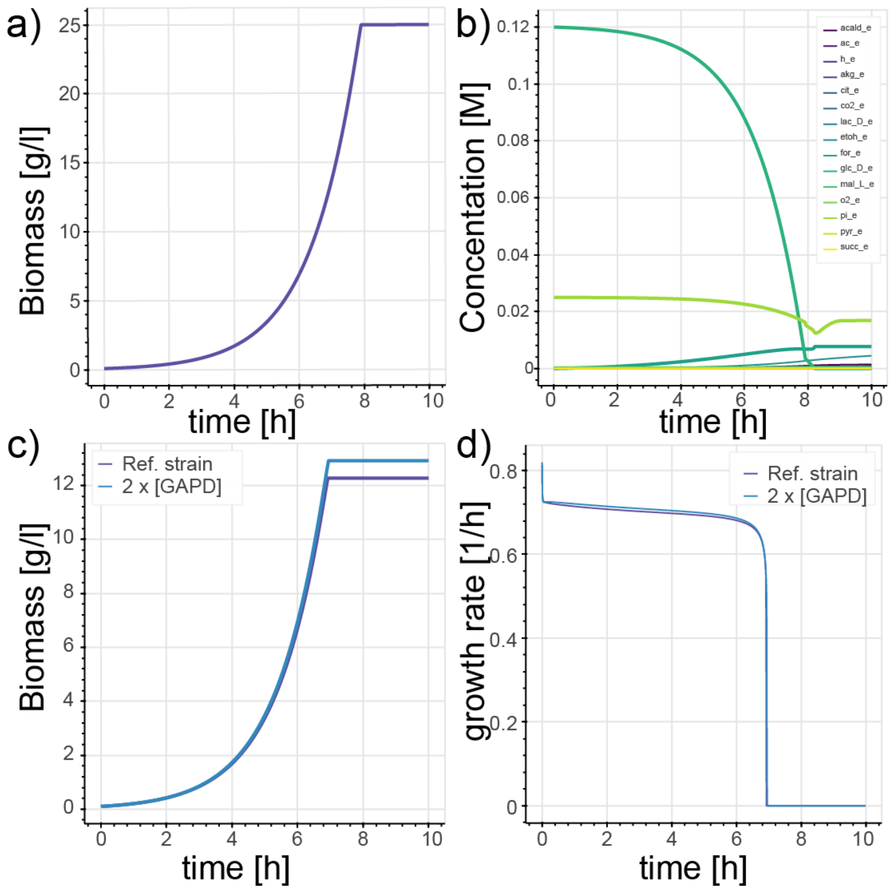


**Figure S5:** a) Biomass formation during the process with a single *E. coli* strain b) Time development of the Medium concentration during the process. c) Biomass formation of a multi-species process with two *E. coli* strains d) Growth rate of the two competing *E. coli* strains.

#### Global sensitivity analysis for metabolic control

Parameter populations estimated using the ORACLE workflow can exhibit large uncertainties in the model behavior. Global sensitivity analysis aids in identifying the parameters contributing most to these uncertainties. Therefore, we implemented a Sobol sensitivity analysis[31] using the Saltelli[32] scheme with efficient resampling according to the ORACLE workflow in SKiMpy. The provided implementation allows decomposing the parametric uncertainty into their main contributors. Such an analysis allows characterizing which model parameters are responsible for uncertainty in the model behavior. In this work, we use SKiMpy to implement a global sensitivity analysis on the metabolic control coefficients, called Global Sensitivity Metabolic Control Analysis (GMCA) in the following[10,11]. The workflow computes which parameters have the most considerable impact on specific metabolic control coefficients using the first order and the total Sobol sensitivity indices[10,11].

We provide a tutorial showing how SKiMpy can be used to apply Global sensitivity analysis to kinetic models. Using a toy model of a branched pathway (based on the upper glycolysis), we apply the GMCA workflow to characterize which model parameters affect the flux control coefficient describing the fold change of the enolase (ENO) flux $v_{ENO}$ for its enzyme activity $[E_{ENO}]$.

$$C_{ENO}^{ENO} = \frac{dln(v_{ENO})}{dln\ ([ENO])}$$

The GMCA workflow uses the ORACLE framework to build a population of locally stable steady-state models. The ORACLE sampling procedure is then used to resample individual parameters efficiently using the Saltelli scheme[32]. In the next step, the control coefficients are calculated using the matrix formulation by Wang et al.[3] Finally, the individual and total sensitivity coefficients are calculated using Saltellis formulation[32,10,11].

In this tutorial, we calculate the individual and total sensitivity of flux control coefficient of the enolase $C_{ENO}^{ENO}$ with respect to the $K_M$s of ENO, phosphate-glycerate mutase (PGM), and phosphate-glycerate kinase (PGK). The results show that for the given fluxes and concentrations, the binding affinity of 2-glycerate mutase (2pg) to ENO, $K_{M,2pg}^{ENO}$, has the strongest individual effect on the control coefficient, followed by the biding of 2pg to PGM, $K_{M,2pg}^{PGM}$ and 3pg to PGK, $K_{M,3pg}^{PGK}$. The results also show that $K_{M,2pg}^{ENO}$ exhibits the smallest total effect indicating that any interaction effects with other parameters are negligible. Interestingly, parameters exhibiting the smallest individual effects, $K_{M,atp}^{PGK}$, $K_{M,adp}^{PGK}$ and, $K_{M,13dpg}^{PGK}$ also have the largest total sensitivity indices suggesting that these parameters exhibit strong interaction effects with other parameters. The results indicate that to reduce the uncertainty of the enolase flux control coefficient its is best advised to reduce the uncertainty of the, $K_{M,2pg}^{ENO}$, $K_{M,2pg}^{PGM}$ and $K_{M,3pg}^{PGK}$ by introducing additional data into the model.

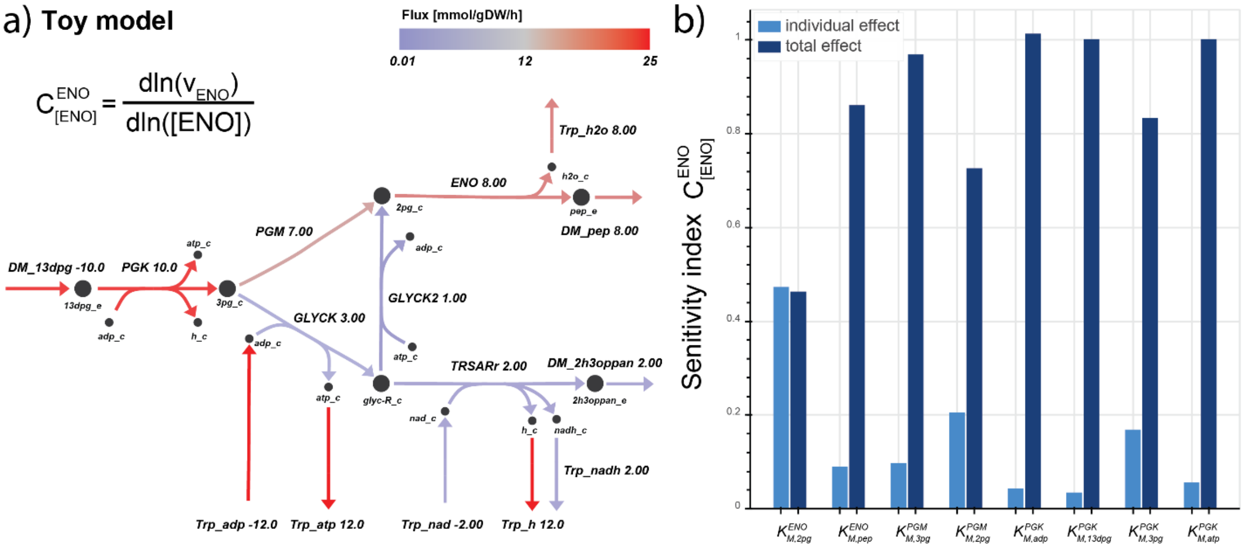


**Figure S6:** Example of a global sensitivity analysis on the control coefficient of enolase (ENO) for a toy model of the central glycolysis in *E. coli*. The model was built around a reference steady-state profile (a) using the ORACLE workflow, when the sensitivity of the flux control coefficient of ENO with respect to its total enzyme concentration was computed for all $K_M$s of ENO, PGM, and PGK (b).

### Supplementary Materials and Methods

#### A metabolic model of the central carbon metabolism of *E. coli*

Aiming to have an educative example to demonstrate methods for the analysis and composition of large-scale kinetic models, we reconstructed a metabolic network representing the central carbon metabolism of *E.coli*, based on the model published by Varma and Palsson[20]. The reconstruction was performed in four steps. First, we identified the 41 reactions from the original model[20] in the *E.coli* genome-scale model (GEM) iJO1366[21]. This resulted in a total of 52 metabolites and 50 reactions belonging to 6 core subsystems (Table S3). Secondly, we include the transport and exchange reactions for the extracellular metabolites, which included carbon sources, inorganics and by-products. In order to have a simple representation of the transport of these compounds, we did not account for the periplasmic compartment and we defined the transport directly between extracellular and cytosol. Thirdly, we integrated in the model the Gibbs free energy of formation for the metabolites and we used thermodynamic flux balance analysis (TFA)[33] to compute the Gibbs free energy of each reactions. This ensures that the directionality of the reactions is in agreement with the laws of thermodynamics. Finally, we wanted to represent the biomass reaction in the model without increasing its complexity. Towards this end, we used lumpGEM[34] to identify the additional reactions from the iJO1366 GEM that are required to produce the biomass building blocks. Then, these reactions are lumped into one general subnetwork which represents the biomass production in the model. The corresponding subnetwork contains 462 reactions and it results in the following lumped reaction:

3665.993 accoa_c + 1144.289 akg_c + 568.838 amp_c + 80049.137 atp_c + 179.126 dhap_c + 391.348 e4p_c + 75.143 f6p_c + 1011.9 for_c + 205 g6p_c + 162.142 glycogenn1_c + 49000.117 h2o_c + 2070.977

nadh_c + 21361.667 nadph_c + 0.352 o2_c + 5586.282 oaa_c + 836.941 pep_c + 568.838 ppi_c + 2935.068 pyr_c + 340.186 q8_c + 965.615 r5p_c + 26.653 ru5p-D_c + 34.286 s7p_c + 545.546 succoa_c + 5.207 ca2_e + 0.235 cbl1_e + 5.207 cl_e + 0.025 cobalt2_e + 0.709 cu2_e + 16.328 fe3_e + 468.428 h_e + 27.841 h2o_e + 195.271 k_e + 8.679 mg2_e + 0.692 mn2_e + 0.144 mobd_e + 11068.711 nh4_e + 0.323 ni2_e + 273.034 so4_e + 0.341 zn2_e <=> 0.235 4crsol_c + 2.602 5drib_c + 630.166 ac_c + 2584.522 acald_c + 80617.975 adp_c + 1.423 amob_c + 2751.173 co2_c + 4211.539 coa_c + 1066.326 fum_c + 50.647 g3p_c + 0.938 glx_c + 54927.291 h_c + 2.83 mththf_c + 2070.977 nad_c + 21361.667 nadp_c + 82306.458 pi_c + 340.186 q8h2_c + 546.85 succ_c + 7.092 5mtr_e + 1.421 btn_e + 5.207 cgly_e + 6.745 glyc_e + 1.423 meoh_e + 1051.595 biomass_e

**Table S3:** Metabolic subsystems in the *E. coli* central carbon metabolism model with the corresponding biotransformation reactions as identified from the iJO1366 based on the model by Varma and Palson.

| **Subsystem** | **Reactions** |
|---|---|
| Glycolysis/Gluconeogenesis | ENO, FBA, FBP, GAPD, PDH, PFK, PGI, PGK, PGM, PYK, TPI, GLCptspp |
| Pentose-phosphate pathway (PPP) | G6PDH2r, GND, PFK_3, PGL, RPE, FBA3, RPI, TALA, TKT1, TKT2 |
| Citric acid cycle (TCA) | ACONTa, ACONTb, AKGDH, CS, FUM, ICDHyr, MDH, SUCOAS |
| Pyruvate Metabolism | ACALD, ACS, ALCD2x, LDH_D, PFL |
| Anaplerotic Reactions | ICL, MALS, ME1, ME2, PPC, PPCK |
| Electron transport chain (ETC) | CYTBO3_4pp, NADH16pp, ATPS4rpp, CYTBDpp, NADH5, SUCDi, NADTRHD, THD2pp, ATPM |

As a result, we generated a model for the central carbon metabolism of *E. coli* containing a total of 96 metabolites, 112 reactions and 121 genes.

#### A kinetic model of the central carbon metabolism of *E. coli*

We next built a kinetic model, based on the reconstructed metabolic model for the carbon metabolism of *E. coli*. As we were interested in understanding the dynamics of the central carbon metabolism of *E.* coli, we simplified the biomass equation to contain only the metabolites that are part of the central carbon metabolism. That is, we neglect 25 metabolites which do not participate in other reactions of the network besides biomass. These included, copper, cobalt, calcium, magnesium, iron, potassium, sulfate, cobalamin, ammonium, nickel, zinc, manganese, chloride, molybdate, biotin, 5-methylthio-D-ribose, glycerol, methanol, cysteyl-glycine, glycogen(n-1), p-cresol, 5''-deoxyribose, S-adenosyl-4-methylthio-2-oxobutanoate, (2R,4S)-2-methyl-2,3,3,4-tetrahydroxytetrahydrofuran. The resulting metabolic model contains 64 metabolites and 65 reactions out of which 16 are transport reactions (Figure S6).

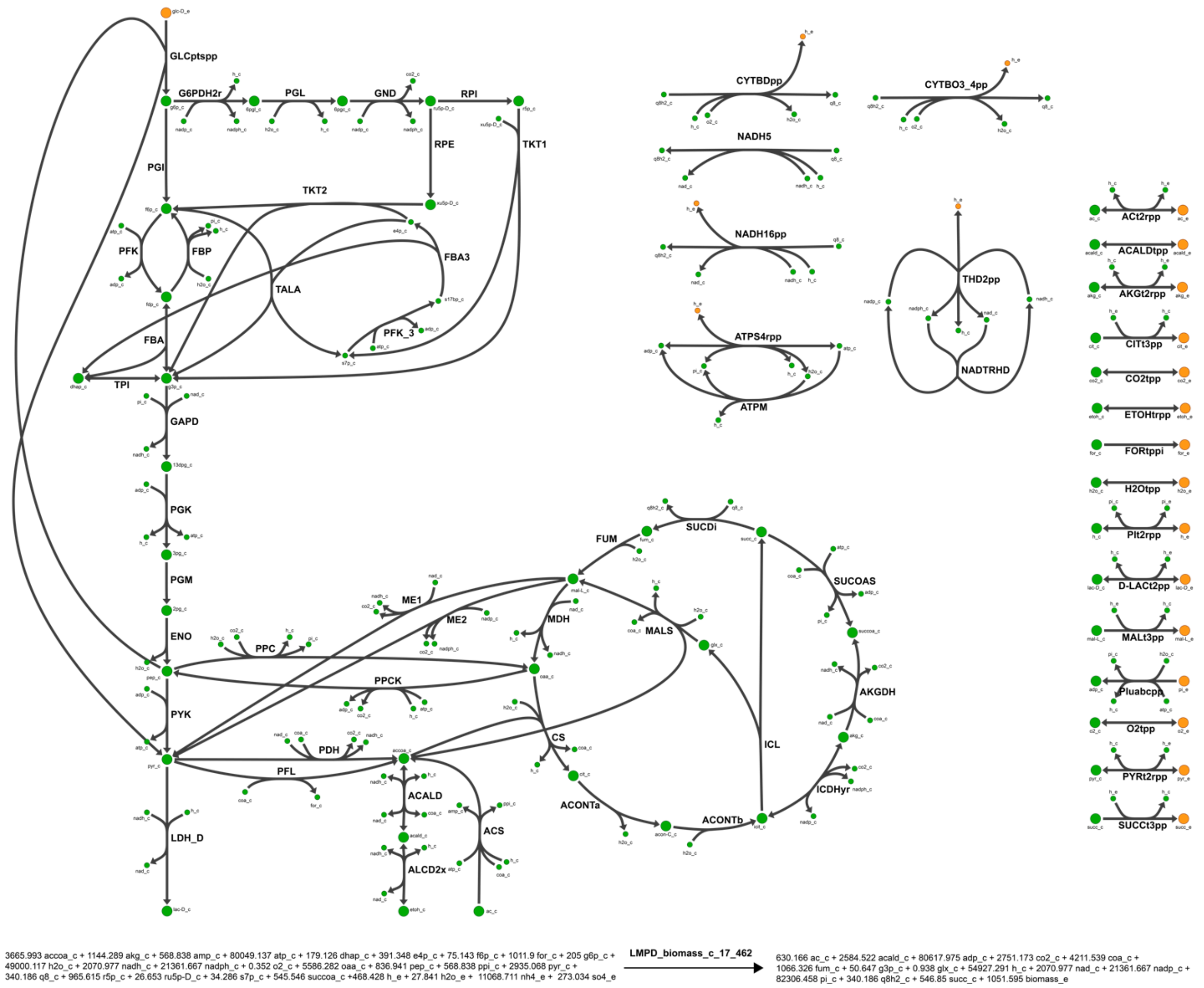


**Figure S7:** Metabolic network of the simplified central carbon metabolism of *E.coli* used as a basis to build a kinetic model. In green the intracellular metabolites, and in orange the extracellular metabolites. The graph was generated with Escher[35].

Next, we used the SKiMpy library (Table S1) to assign reaction kinetics accordingly to the stoichiometry of each reaction. Additionally, in this case, we defined protons as small molecules, i.e., protons are not part of the mechanism stoichiometry but are considered as small molecule modifiers (Table S2) that change the forward and backward reaction rate as a function of the small molecule concentrations and account for their stoichiometric balance. As a result of assigning kinetic mechanisms to the 65 reactions in the model, we identified that 23 reactions are described with a reversible Michaelis-Menten mechanism, four reactions have Uni-Bi reversible hill, one reaction has Bi-Uni reversible hill, 23 reactions have generalized reversible hill (19 with a Bi-Bi stoichiometry and 4 with a Ter-Ter stoichiometry), eight were assigned convenience kinetics mechanisms and six reactions (Table S4).

**Table S4:** Kinetic mechanisms in the central carbon metabolism of *E. coli* kinetic model.

| **Mechanism** | **Stoichiometry** | **Reactions** |
|---|---|---|
| Reversible Michaelis-Menten | $S_1 \rightleftharpoons P_1$ | ACALDtpp, ACONTa, ACONTb, ACt2rpp, AKGt2rpp, CITt3pp, CO2tpp, D-LACt2pp, ENO, ETOHtrpp, FORtppi, FUM, MALt3pp, O2tpp, PGI, PGL, PGM, PIt2rpp, PYRt2rpp, RPE, RPI, SUCCt3pp, TPI (23) |
| Uni-Bi Reversible Hill | $S_1 \rightleftharpoons P_1 + P_2$ | FBA, FBA3, FBP, ICL (4) |
| Bi-Uni Reversible Hill | $S_1 + S_2 \rightleftharpoons P_1$ | ATPS4rpp (1) |
| Generalized Reversible Hill | $S_1 + S_2 \rightleftharpoons P_1 + P_2$ | ALCD2x, CS, G6PDH2r, GLCptspp, LDH_D, MALS, MDH, NADTRHD, PFK, PFK_3, PFL, PGK, PPC, PYK, SUCDi, TALA, THD2pp, TKT1, TKT2 (19) |
| | $S_1 + S_2 + S_3 \rightleftharpoons P_1 + P_2 + P_3$ | ACS, AKGDH, PDH, SUCOAS (4) |
| Convenience Kinetics | $S_1 + S_2 + S_3 \rightleftharpoons P_1 + P_2$ | ACALD, GAPD (2) |
| | $S_1 + S_2 \rightleftharpoons P_1 + P_2 + P_3$ | GND, ICDHyr, ME1, ME2, PPCK (5) |
| | $S_1 + S_2 \rightleftharpoons 2P_1 + P_2$ | PIuabcpp (1) |
| Irreversible Michaelis-Menten | $S_1 \rightleftharpoons P_1 + P_2$ | ATPM (1) |
| | $S_1 + 2S_2 \rightleftharpoons 2P_1$ | CYTBDpp, CYTBO3_4pp (2) |
| | $S_1 + S_2 \rightleftharpoons P_1 + P_2$ | NADH16pp, NADH5 (2) |
| | Complex* | LMPD_biomass_c_17_462 (1) |

Then, we formulate the ordinary differential equations (ODEs) describing the mass balances of the system. Out of the 64 metabolites in the metabolic model, 49 have an ODE for their mass balance, and 15 metabolites are considered as boundary metabolites with known concentrations.

These stoichiometric, thermodynamic, and kinetic models are used as examples in several tutorials to showcase the computational and methodical capabilities of SKiMpy.

#### Steady-State consistent model parameters (ORACLE workflow)

In this section, we demonstrate how SKiMpy can be used to build large-scale Kinetic models using the ORACLE workflow. In many cases, the available experimental data is insufficient to parameterize large-scale kinetic models. The ORACLE workflow addresses this problem by sampling unknown parameters. Therefore, the ORACLE workflow uses the thermodynamic-based flux analysis to compute a thermodynamically feasible set of steady-state fluxes and concentrations. These steady-state fluxes and concentrations serve then as a basis to sample parameters that are consistent with the steady-state. Specifically, all parameters affecting the saturation of an enzymatic reaction are sampled with respect to a reference concentration, including Michaelis-Menten constants and inhibitory constants. Therefore, each constant $K_M^{ij}$ is assigned to a respective steady-state concentration $[S_i]$. This parameter concentration pair is then treated as if it contributed to the saturation of the enzyme in the form of the hyperbolic function known from the irreversible Michaelis-Menten kinetics[3]:

$$\sigma_{ij} = \frac{[S_i]/K_M^{ij}}{1 + [S_i]/K_M^{ij}}.$$

This formulation allows replacing the unbounded sampling of Michaelis-Menten constants with a simple sampling on the $[0,1]$ interval[3]. Back insertion of the kinetic and thermodynamic parameters, i.e. $K_M^{ij}$ and $K_{eq}^{j}$, in the rate expression allows calculating the respective maximal enzyme velocities $V_{max}^{j}$ based on the corresponding steady-state flux $v_j$. The resulting parameter sets are then consistent with the assumed steady-state profile (Figure S8).

In the next step, we follow the formulation by Wang et al.[3] to calculate the Jacobian of the dynamic system for every parameter sample. In large-scale metabolic models, linear dependencies in the stoichiometric matrix result in pools of metabolites that remain constant[36]. Such systems require computing the reduced stoichiometric matrix $\boldsymbol{N_R}$ to compute of the Jacobian[8]. This reduced matrix contains only the linear independent rows of the original stoichiometric matrix $\boldsymbol{N}$. Multiple algorithms have been proposed to identify such pools[37–39]. For large-scale metabolic models, we recommend using the MILP formulation presented by Nicolaev et al.[14]. A MATLAB implementation of this method can be found at: https://github.com/EPFL-LCSB/matMCP.

As mentioned above, the eigenvalues of Jacobian define the characteristics of the system close to the reference steady-state:

$$y_i(t) = e^{\lambda_i t}$$

The reciprocal of the real part of the Jacobian's eigenvalues provide a characteristic time constant of the linearized system. Moreover, a Jacobian whose largest eigenvalue has a positive real part will yield unstable models. Therefore, each sampled parameter set is evaluated with respect to its local stability.

Also, eigenvalues with a negative real part too close to 0 will yield slow dynamics, potentially slower than metabolism. As a result, it is necessary to filter the parameter sets to discard (i) unstable models and (ii) models with slow dynamics.

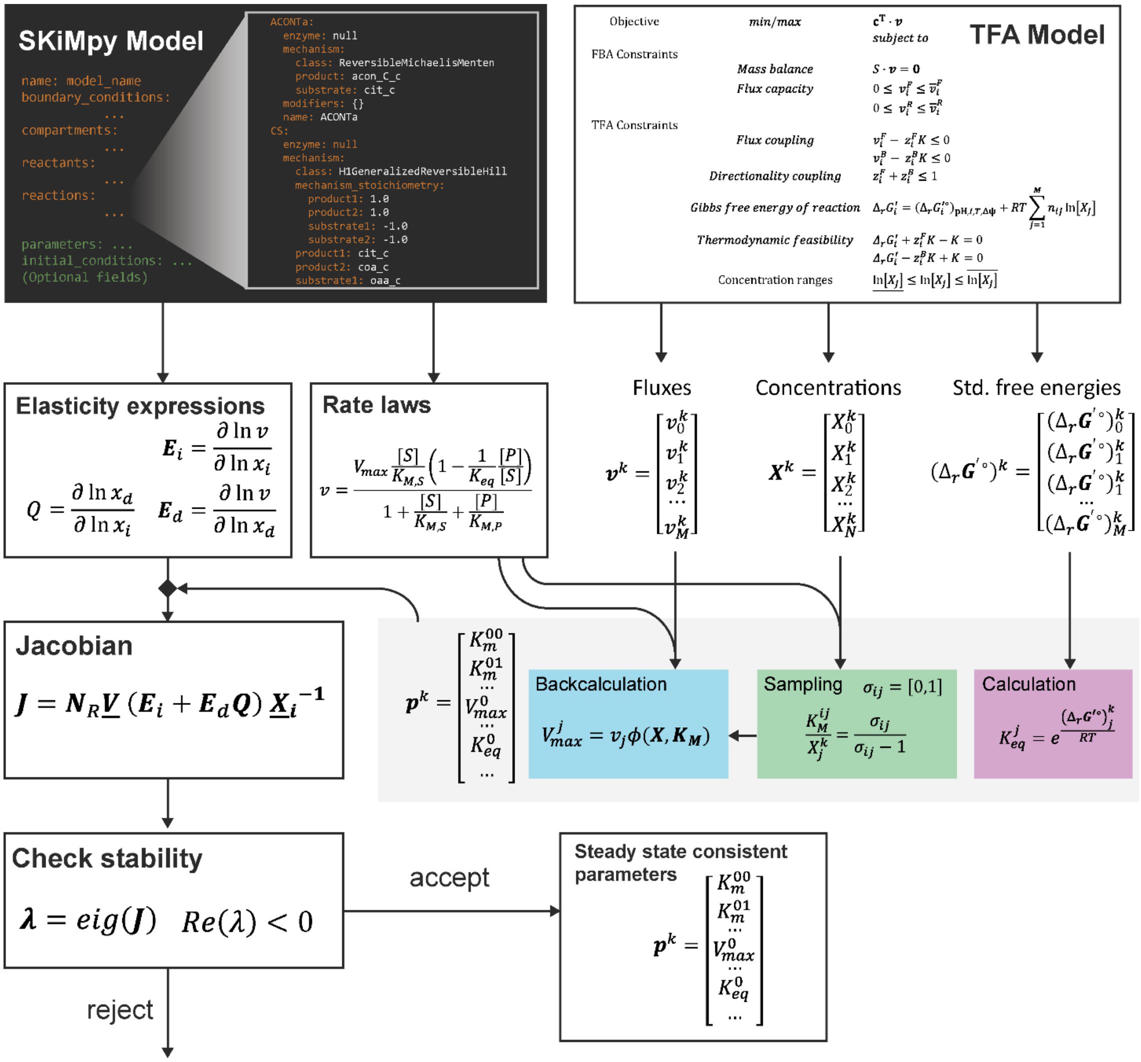


**Figure S8:** Workflow to sample stable steady-state consistent parameter sets for large-scale kinetic models (ORACLE workflow).

(15) Meurer, A.; Smith, C. P.; Paprocki, M.; Čertík, O.; Kirpichev, S. B.; Rocklin, M.; Kumar, Am.; Ivanov, S.; Moore, J. K.; Singh, S.; Rathnayake, T.; Vig, S.; Granger, B. E.; Muller, R. P.; Bonazzi, F.; Gupta,

H.; Vats, S.; Johansson, F.; Pedregosa, F.; Curry, M. J.; Terrel, A. R.; Roučka, Š.; Saboo, A.; Fernando, I.; Kulal, S.; Cimrman, R.; Scopatz, A. SymPy: Symbolic Computing in Python. *PeerJ Comput. Sci.* **2017**, *3*, e103. https://doi.org/10.7717/peerj-cs.103.

(23) Alberts, B. *Essential Cell Biology*, 4th ed.; Garland Science: New York, 2014.

(24) Salvy, P.; Hatzimanikatis, V. The ETFL Formulation Allows Multi-Omics Integration in Thermodynamics-Compliant Metabolism and Expression Models. *Nat Commun* **2020**, *11* (1), 1–17. https://doi.org/10.1038/s41467-019-13818-7.

(25) Schuster, S.; Schuster, R. A Generalization of Wegscheider's Condition. Implications for Properties of Steady States and for Quasi-Steady-State Approximation. *J Math Chem* **1989**, *3* (1), 25–42. https://doi.org/10.1007/BF01171883.

(26) Hanekom, A. J. Generic Kinetic Equations for Modelling Multisubstrate Reactions in Computational Systems Biology. Master Thesis, Stellenbosch : University of Stellenbosch, 2006.
